# Establishment of a social conditioned place preference paradigm for the study of social reward in female mice

**DOI:** 10.1101/2022.03.31.486566

**Authors:** Zofia Harda, Magdalena Chrószcz, Klaudia Misiołek, Marta Klimczak, Łukasz Szumiec, Maria Kaczmarczyk-Jarosz, Jan Rodriguez Parkitna

## Abstract

Social interactions can be and often are rewarding. The effect of social contact strongly depends on circumstances, and the reward may be driven by varied motivational processes, ranging from parental or affiliative behaviors to investigation or aggression. Reward associated with nonreproductive interactions in rodents is measured using the social conditioned place preference (sCPP) paradigm, where a change in preference for an initially neutral context confirms reinforcing effects of social contact. Here, we revised the sCPP method and reexamined social reward in adult female mice. Contrary to earlier studies, we found that robust rewarding effects of social contact could be detected in adult (14-week-old) female C57BL/6 mice when the sCPP task was refined to remove confounding factors. Strikingly, the rewarding effects of social interaction were only observed among female siblings who remained together from birth. Contact with same-age nonsiblings was not rewarding even after 8 weeks of cohousing. Other factors critical for the social reward effect in the sCPP paradigm included the number of conditioning sessions and the inherent preference for contextual cues. Thus, we show that social interaction is rewarding in adult female mice, but this effect strictly depends on the familiarity of the interaction partners. Furthermore, by identifying confounding factors, we provide a behavioral model to study the mechanisms underlying the rewarding effects of nonreproductive social interaction in adult mice.

## Introduction

The rewarding effects of social contact are the proximate cause for all forms of interaction with conspecifics. The direct method to model the rewarding effects of nonreproductive interaction in rodents is the social conditioned place preference (sCPP) paradigm, where social contact causes an increase in the time spent in a previously neutral context^1^. In the test, animals are exposed to a set of environmental cues when placed in isolation and a second, different set of cues when housed in a group. Accordingly, when animals are tested for preference between the two sets of cues, an increase in the preference for the social context is used as a measure of social reward. sCPP directly measures the reinforcing effect of social contact and is based on the same construct as the classic conditioned place preference test that is used extensively to test the rewarding effects of drugs. The sCPP protocol, with various modifications, has been instrumental in a series of studies showing the roles of serotonergic and oxytocin-dependent signaling in neuronal plasticity underlying social reward^2–6^. Reported results show that rewarding effects of social contact could be robustly observed in juvenile animals; however, the effect of social interaction is diminished or absent in early adulthood at the age of 8 weeks in male mice and at 11 weeks in females^5^. These results could be considered unexpected, as a lack of sCPP in adult mice would be at odds with other indices of social interaction preference in adult mice, such as the anxiogenic effects of isolation^7^ or communal nesting female behavior^8^.

A method suitable for studying social reward in adult mice would resolve two basic issues. Primarily, the rewarding effects of social contact in mature adults likely differ in underlying mechanisms from those observed in juveniles, i.e., if rewarding effects of social contact without reproductive context in adult model animals can be reliably demonstrated. Moreover, while models of impaired social behavior in juveniles have obvious translational value, neuropsychiatric disorders affecting adults also frequently involve altered social behavior. Second, using adults as model animals facilitates the application of genetic tools to study neuronal signaling involved in social interaction. Commonly used methods to assess neural circuit activity require periods of weeks to achieve optimal transgene activity; thus, they are not well suited for the study of juvenile mice. Furthermore, when using genetically modified rodents, breeding large cohorts with identical ages is very difficult in practice. This, combined with the relatively high number of animals required to achieve an adequate test power, has made application of this task in mutant mice impractical.

Here, we present a revised sCPP task that reliably measures social reward in adult C57BL/6 mice. We focused exclusively on females, since observations of Mus musculus behavior in natural and semi-natural environments suggest that female mice may form small groups and display communal nesting and nursing behavior^8^. Conversely, interactions between male mice are rare and usually antagonistic^9^. Our results indicate that social contact is rewarding in mature adult female C57BL/6 mice; however, the effect is dependent on the context, conditioning length, and familiarity.

## Results

### Neutral conditioning contexts

The sCPP paradigm is based on Pavlovian conditioning, where a context (i.e., a neutral stimulus) acquires the capacity to elicit a response if paired with a reward (i.e., social interaction). Therefore, the first step was to select a set of contexts not associated with inherent preferences in mature adult female C57BL/6 mice, which would permit an unbiased assignment of the initial bedding context. A schematic representation of the testing apparatus is shown in Figure 1A. Preference for four pairs of contexts was assessed, each context consisting of a distinct type of bedding and a wooden gnawing block. Bedding types differed in odor, texture and color, and gnawing blocks differed in shape and size. Before the experiment, a separate type of bedding and gnawing block was used in the home cages, so the animals had no previous exposure to the conditioning contexts. Preference was tested in 10- to 17-week-old female mice in a 30-minute session, during which mice could freely move between the two cage compartments (Figure 1A). As summarized in Figure 1B, mice had a significant inherent preference for one of the context types in sets 3 and 4 (one-sample t_22_ = 4.163, p < 0.001 and t_6_ = 9, p < 0.001, respectively, see Table S1 for extended data), and these sets were not used in further experiments. In the case of set 2, while no significant preference between contexts was detected despite the apparent trend (t_15_ = 1.965, p = 0.068), mice were observed to be eating one of the bedding types, and thus, set 2 was excluded as well. Only in one case, set 1, was no inherent preference observed (t_113_ = 0.119, p = 0.906, the data include initial preference from all results shown in the manuscript). No significant differences in the distance traveled between the different sets were observed (Figure 1C, one-way ANOVA F_3,156_ = 2.615, p = 0.053). Based on these results, set 1 (beech type 1 + cellulose) was selected for all further experiments as being completely neutral during initial exposure.

**Figure 1.**
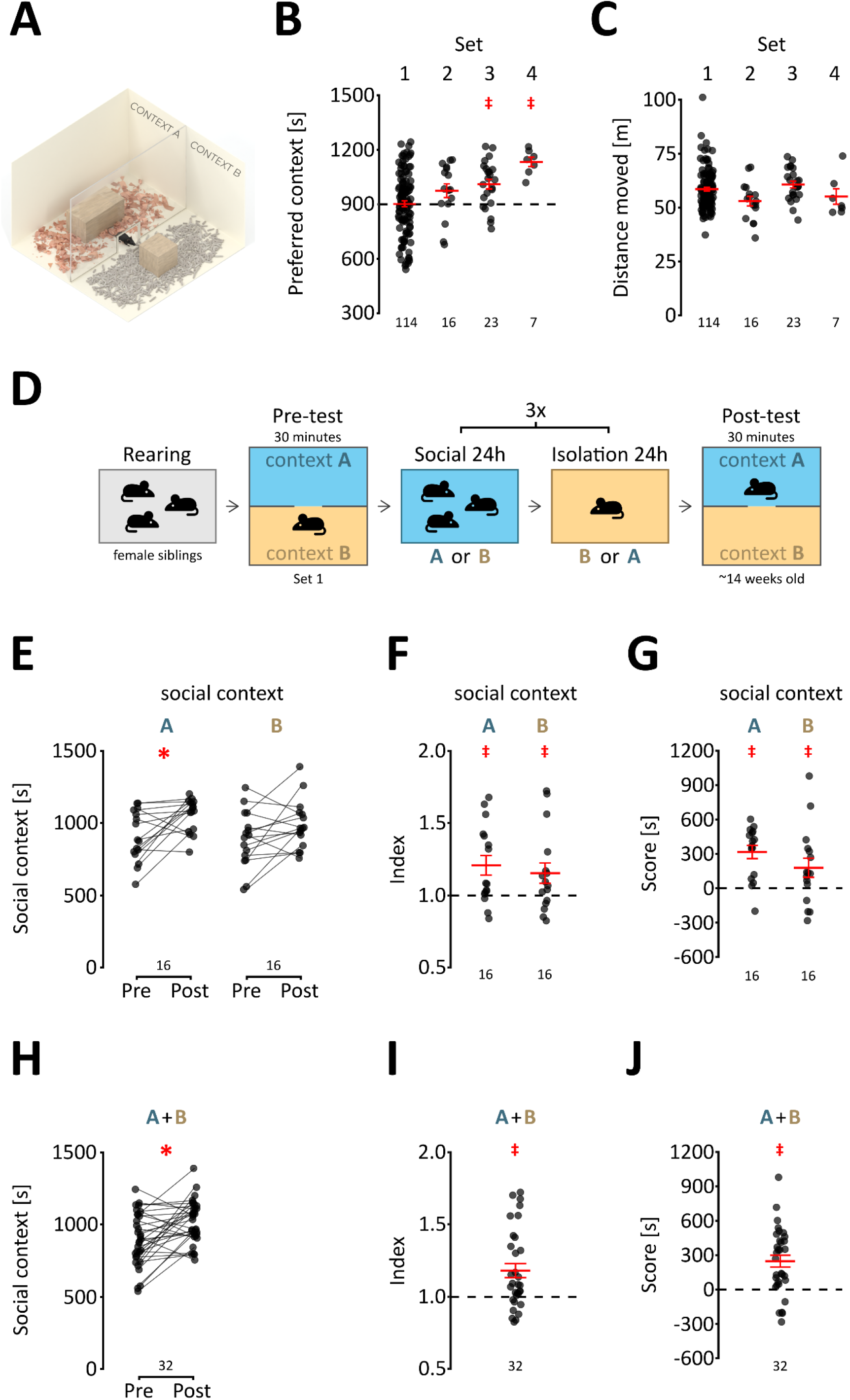
Social reward in adult mice. (A) A schematic representation of the conditioning cage. The cage was a 30×30×30-cm cube. The partition is a transparent Plexiglas plate with an opening at the base that allows the animal to cross freely. (B) Inherent preference of conditioning contexts. Each point represents the time spent in the preferred compartment (i.e., having greater average preference) by a single animal. The total time of the test was 1800 s (30 min), and the dashed line represents equal time spent in both compartments (900 s). The sets of bedding used for conditioning are indicated above the graph, complete information on bedding types is provided in supplementary Tables S1 and S4. The numbers of animals tested in each set are shown below the graph. The mean and s.e.m. are shown in red, and significant preference is marked with a red “‡” (p < 0.05, one-sample t test vs. 900 s). (C) Distance traveled. The graph shows the distance traveled by each animal during the 30-minute test. Sets are indicated above, and the group sizes are shown below the graph. The mean and s.e.m. values are shown in red. (D) Schematic representation of the conditioning procedure. The bedding sets are listed in supplementary Table S1. (E) Time spent in context associated with social interaction during the pre- and post-test. For each animal, the times spent in the social context during the pre- and post-tests are joined by a line. A significant increase from the pre-to post-test is marked with a red “*” (p < 0.05, Bonferroni corrected t test). Group sizes are indicated below the graph. (F) Preference index. The values represent the ratio between the time spent in the social context during the post-test to the corresponding time during the pre-test. Group sizes are shown below the graph, and the mean and s.e.m. values are shown in red. A significantly greater than 1 mean index value is shown as a red “‡” (p < 0.05, one-sample t test vs. 1). (G) Preference score. The values represent the difference in time spent in the social context and isolation context during the post-test. Group sizes are shown below the graph, and the mean and s.e.m. values are shown in red. A significant mean preference score is shown as a red “‡” (p < 0.05, one-sample t test vs. 0). (H, I, J) The change in time spent in the first context between pre- and post-test, the index and the score, respectively, data pooled for contexts A and B. The mean and s.e.m. values are shown in red, where appropriate. Significant effects are shown with a red “*” or “‡” (p < 0.05, two- or one-sample t test, respectively).

### Social reward in adult female mice

Next, we assessed whether social contact with same-sex, same-age siblings was rewarding for adult female C57BL/6 mice (14 weeks old, see Table S3). The test followed a similar design as previously used in the case of juvenile mice^1,3,10^. Briefly, mice underwent 6 days of conditioning, during which they were moved between a social cage with siblings and single housing with the respective contexts every 24 hours (Figure 1D). The assignment of beech (A) and cellulose (B) contexts was randomized between litters to preserve a balanced and unbiased design. An increase in time spent in the context associated with social interaction (the reward stimulus) was interpreted as a conditioned response and thus proves that the stimulus acted as a reward. The effect of conditioning was assessed either by the relative change in preference between pre- and post-test (“Pre Post” and “Index”) or the preference for the social-conditioned context during the post-test (“Score”).

Adult female mice had a significantly increased preference for the social context from the pre-test to the post-test when A (beech) was paired with social contact (Figure 1E, *context* F_1,30_ = 0.871, p = 0.358, *pre-post* F_1,30_ = 12.526 p = 0.001; *interaction* F_1,30_ = 0.509 p = 0.481; *post hoc* Bonferroni t test context A and B: t_15_ = 3.021 p = 0.017 and t_15_ = 1.989 p = 0.131, respectively), which was also evident in the preference index (Figure 1F, one-sample t test context A and B: t_15_ = 3.069 p = 0.008 and t_15_ = 2.188 p = 0.045, respectively). Likewise, mice had a significant preference for the social context during the post-test, irrespective of the type of bedding (Figure 1G, one-sample t test context A and B: t_15_ = 5.492 p < 0.001, t_15_ = 2.136 p = 0.050, respectively). In the case of context A, 15 out of 16 animals (94%) had a preference for the social-conditioned context, and the same was the case for 12 out of 16 (75%) mice for context B. There were no significant effects of the type of context (A or B). When the data from both contexts were pooled, the social reward effect was significant in all three measures: the increase in time spent in the context (Figure 1H&I, t_31_ = 3.568 p = 0.001 and t_31_ = 3.747 p < 0.001, respectively) and the preference during the post-test (Figure 1J, t_31_ = 4.808 p < 0.001). Together, these results show that adult (14 weeks old) female C57BL/6 mice acquired a preference for the initially neutral, social-conditioned context and thus demonstrate that social contact with sibling females was rewarding.

### Social contact independent effects

Next, we considered the possibility that the context itself (i.e., the type of bedding) could act as a reward. Speculatively, while there was no initial preference between contexts, one of the types of bedding would become preferred during conditioning. Additionally, it could have been argued that mice showed a preference for the context they were deprived of during the previous 24 h, irrespective of social interaction. To test these possibilities, we conducted an experiment following the same procedure as before, except that the mice remained with siblings in both contexts (Figure 2A). The results from these experiments are shown separately for the two starting contexts. When the mice were first exposed to context A, an increase in time spent in the A context was observed, although both A and B were paired with social contact (“noncond”, Figure 2B). There were no significant differences between time spent in context A, when only this context was paired with social interactions compared to when both contexts were associated with social contact (*cond* vs. *noncond* F_1,26_ = 0.273, p = 0.606, *pre-post* F_1,26_ = 12.92 p = 0.001, *interaction* F_1,26_ = 0.00 p = 0.995, *post hoc* Bonferroni t test for the “noncond” Group t_11_ = 2.094, p = 0.120). In line with this, there were no significant differences in the average index and score between the conditioned and nonconditioned groups when the animals were first exposed to context A (Figure 2C&D, t_26_ = 0.060 p = 0.953 and t_26_ = −0.452 p = 0.655, respectively). The index and score in the nonconditioned group were higher than the chance level (t_11_ = 2.4622 p = 0.032 and t_11_ = 3.332 p = 0.007, respectively). Conversely, in the case where the first exposure was to context B, there was a significant interaction of pre vs. post difference and group i.e., conditioned vs. nonconditioned mice (Figure 2E, *conditioning* F_1,23_ = 3.306, p = 0.082, *pre-post* F_1,23_ = 0.583 p = 0.453, *interaction* F_1,23_ = 6.404 p = 0.0187, *post hoc* Bonferroni t test for the “noncond” Group t_8_ = −1.839 p = 0.206) and significantly lower index and score compared to the normal procedure (Figure 2F&G, t_23_ = −2.243 p = 0.035 and t_23_ = −3.308 p = 0.003, respectively). The difference between the “cond” and “noncond” groups would have been in line with expectations, as the “noncond” group had not develop preference for any of the contexts. Nevertheless, the score value in the “noncond” group was significantly lower than the chance level (t_8_ = −2.832 p = 0.022), thus again pointing to a development of context A preference. These data show that while context A is initially neutral, it becomes preferred following repeated exposure, irrespective of social conditioning. However, this effect can be controlled by balancing the number of animals assigned to the two starting contexts. Imbalances occurred despite planning equally sized groups due to a greater than 70% initial preference for one of the contexts and variability in litter sizes. To make the groups equal in size, we randomly trimmed the larger group using an R script. When equal numbers of animals for both starting contexts were combined, no conditioning effects were observed in the nonconditioned group in the pre-vs. post-test difference (Figure 2H, t_17_ = 0.514 p = 0.614), index (Figure 2 I, t_17_ = 1.026 p = 0.320) or score (t_17_ = 0.323 p = 0.614). A comparison of the data before and after trimming, with the results corresponding to each context separately are shown in the Supplemental Figure S1. The pairwise comparisons of contexts again reveal an effect related to the bedding, and show that the reduction of the larger group did not appreciably alter the change in time spent from pre-to post-test, preference index or preference score. Taken together, although we observed context-type conditioning, the effect could be separated from the social contact reward. For all further experiments, the experiments were randomly ‘trimmed’ in the same fashion to balance the contexts.

**Figure 2.**
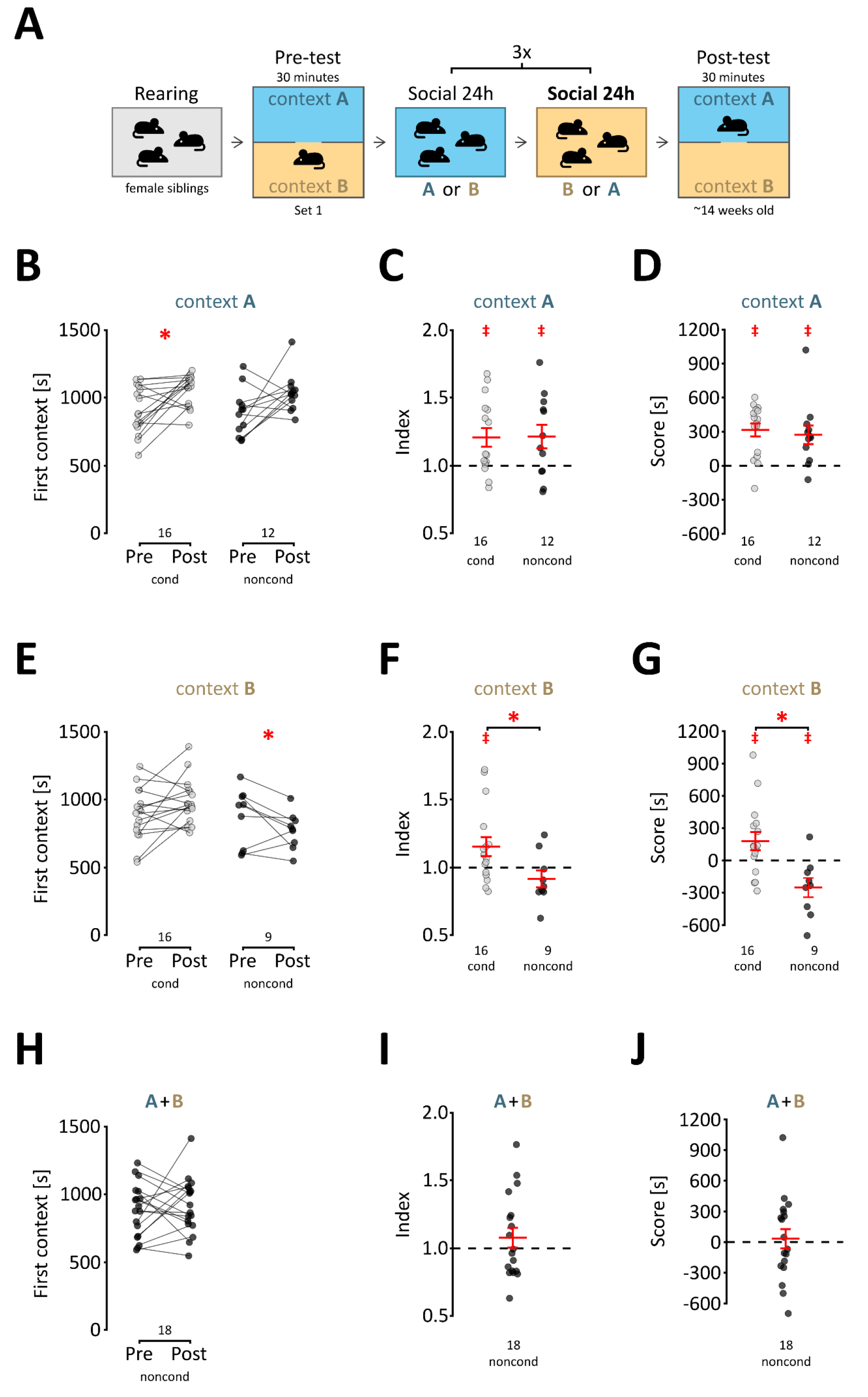
Social interaction-independent effects. (A) Schematic representation of the procedure; both contexts are associated with social interactions. (B) Time spent in context A during the pre- and post-test. For each animal, the times spent in the first context during pre- and post-tests are joined with a line. “Noncond” represents animals that remained with siblings in both contexts. The data in the “cond” represent normal conditioning and are shown for comparison (the same data are shown in 1E). For the “cond” group, the “first context” equals the social context. A significant increase from the pre- to post-test is marked with a red “*” (p < 0.05, two-sample t test). Group sizes are indicated below the graph. (C) Preference indices for mice first exposed to context A. The values represent the ratio between the time spent in the social context during the post-test to the corresponding time during the pre-test. “Noncond” represents animals that remained with siblings in both contexts. The data in the “cond” represent normal conditioning (same as 1F) and are shown for comparison. Group sizes are shown below the graph, and the mean and s.e.m. values are shown in red. A significantly greater than 1 mean index value is shown as a red “‡” (p < 0.05, one-sample t test vs. 1). (D) Preference scores for mice first exposed to context A. The values represent the difference in time spent in the social context and isolation context during the post-test. “Noncond” represents animals that remained with siblings in both contexts. The data in the “cond” represent normal conditioning (same as 1G) and are shown for comparison. Group sizes are shown below the graph, and the mean and s.e.m. values are shown in red. A significant mean preference score is shown as a red “‡” (p < 0.05, one-sample t test vs. 0). (E, F and G) These graphs are the equivalents of B, C and D, respectively, but for mice first exposed to context B. (H, I and J) The change in time spent in the first context between pre- and post-test, the index and the score, respectively. These graphs show merged data for both first contexts for the “noncond” group.

### Conditioning length and familiarity

Previously, it was reported that no significant rewarding effects of social contact were observed in C57BL/6J female mice that were older than 11 weeks at post-test^5^. However, these data were based on a protocol with only one 24-h social context conditioning period, followed by one 24-h isolation period. Indeed, when we shortened the conditioning to 2 days, 14-week-old adult C57BL/6 female mice from the IP PAS stock (see Table S3) showed no significant increase in social context preference from pre-to post-test (Figure 3A, t_23_ = 0.805 p = 0.430), and there were no significant effects on the index and score (Figure 3 B&C, t_23_ = 0.8853 p = 0.385 and t_23_ = 0.773 p = 0.448, respectively). Conversely, when the 8-week-old female mice underwent conditioning in the 2-day protocol, a significant increase in preference from pre-to post-test was observed (Figure 3D, t_19_ = 3.316 p = 0.004), and the score and index were significantly above the chance level (Figure 3E&F, t_19_ = 3.317 p = 0.004 and t_19_ = 3.795 p = 0.001, respectively). These results replicate previously reported data^5^ and show that the length of the conditioning strongly and differentially affects the sCPP in mice depending on age.

**Figure 3.**
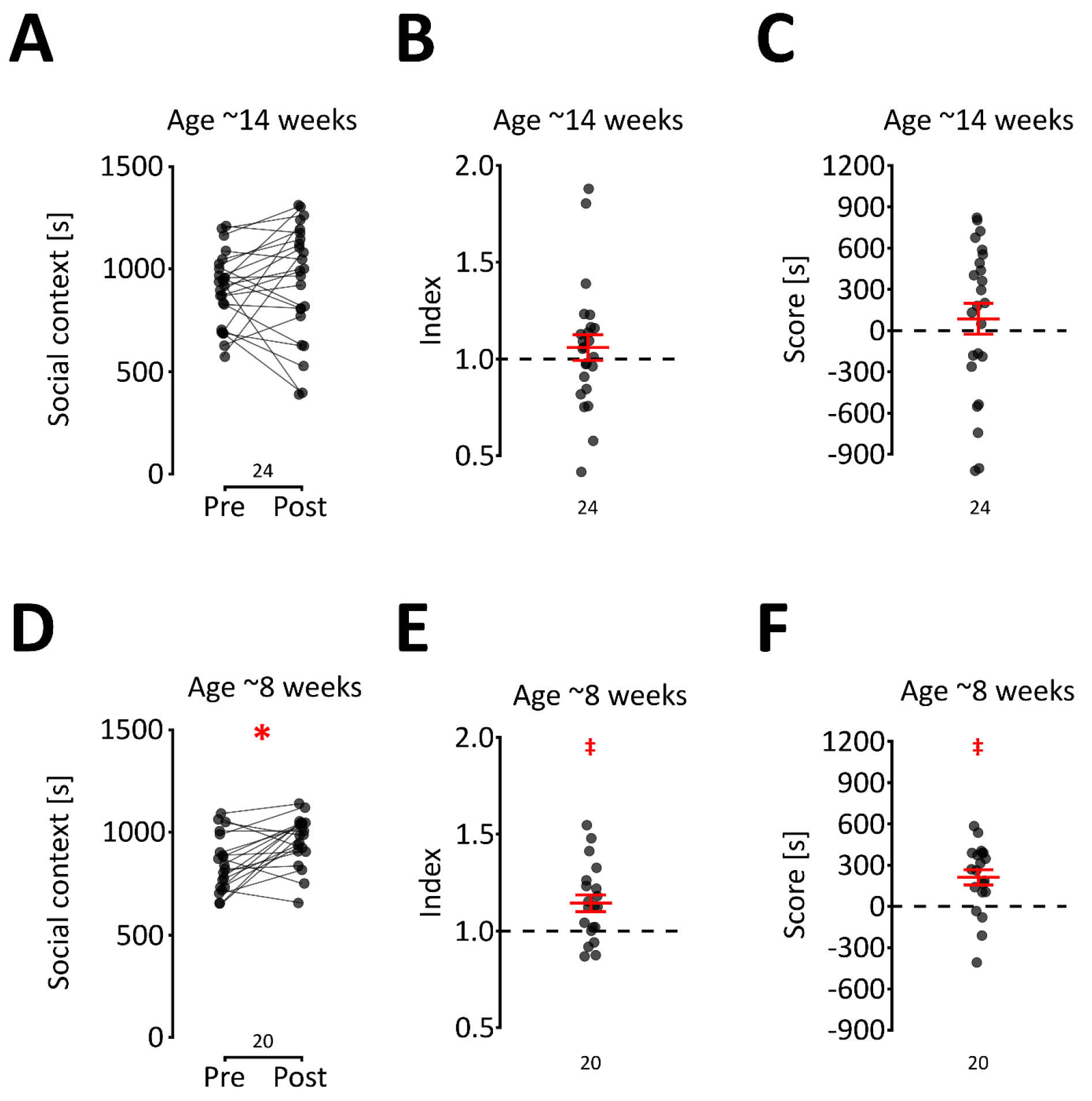
The effect of the time length on conditioning on social reward. (A, B and C) The change in time spent in the first context between pre- and post-test, the index and the score, respectively, for 8-week-old mice after the 2-day conditioning procedure. The mean and s.e.m. values are shown in red in Panels B & C. (D, E and F) The change in time spent in the first context between pre- and post-test, the index and the score, respectively, for 8-week-old mice after the 2-day conditioning procedure. The mean and s.e.m. values are shown in red in Panels E & F. Significant effects are shown with a red “*” or “‡” (p < 0.05, two- or one-sample t test, respectively).

Social behaviors are strongly influenced by familiarity; therefore, in the final experiment, we assessed sCPP in two cohorts that were not siblings and were housed together for a specific period of time before the test. The first cohort comprised mice that were combined after weaning (i.e., ~4 weeks of age), while the second cohort consisted of mice that were brought together one day before the pre-test (Figure 4A). In both experiments, mice were brought together to form groups of 3 to 5 in such a way that every animal in the group was from a different litter. The conditioning procedure was started when the mice reached at least 12 weeks of age. Strikingly, we found no effect of conditioning on context preference in mice that had been housed together since weaning (Figure 4B, C and D, t_17_ = −0.095 p = 0.926, t_17_ = −0.081 p = 0.936 and t_17_ = 0.556 p = 0.586, respectively). The same was the case in animals that were brought together a day before the pre-test (Figure 4E, F and G, t_17_ = 1.249 p = 0.229, t_17_ = 1.748 p = 0.098 and t_17_ = 0.98911 p = 0.337, respectively). Thus, we did not observe rewarding effects of social interaction in nonsibling mice, even when they were housed together for a period of ~8 weeks, although it remains uncertain whether the critical factor was the age when the animals were brought together or their relatedness. Additionally, the results from this experiment indicate that the preference for the social context is not driven by aversion to the isolation context, barring the unlikely case where interaction with nonsiblings and isolation are equally aversive. If it had been the aversion to the isolation context causing the preference for the social context in the post-test, then preference for social context would have been observed in the case of nonsiblings, which was not the case.

**Figure 4.**
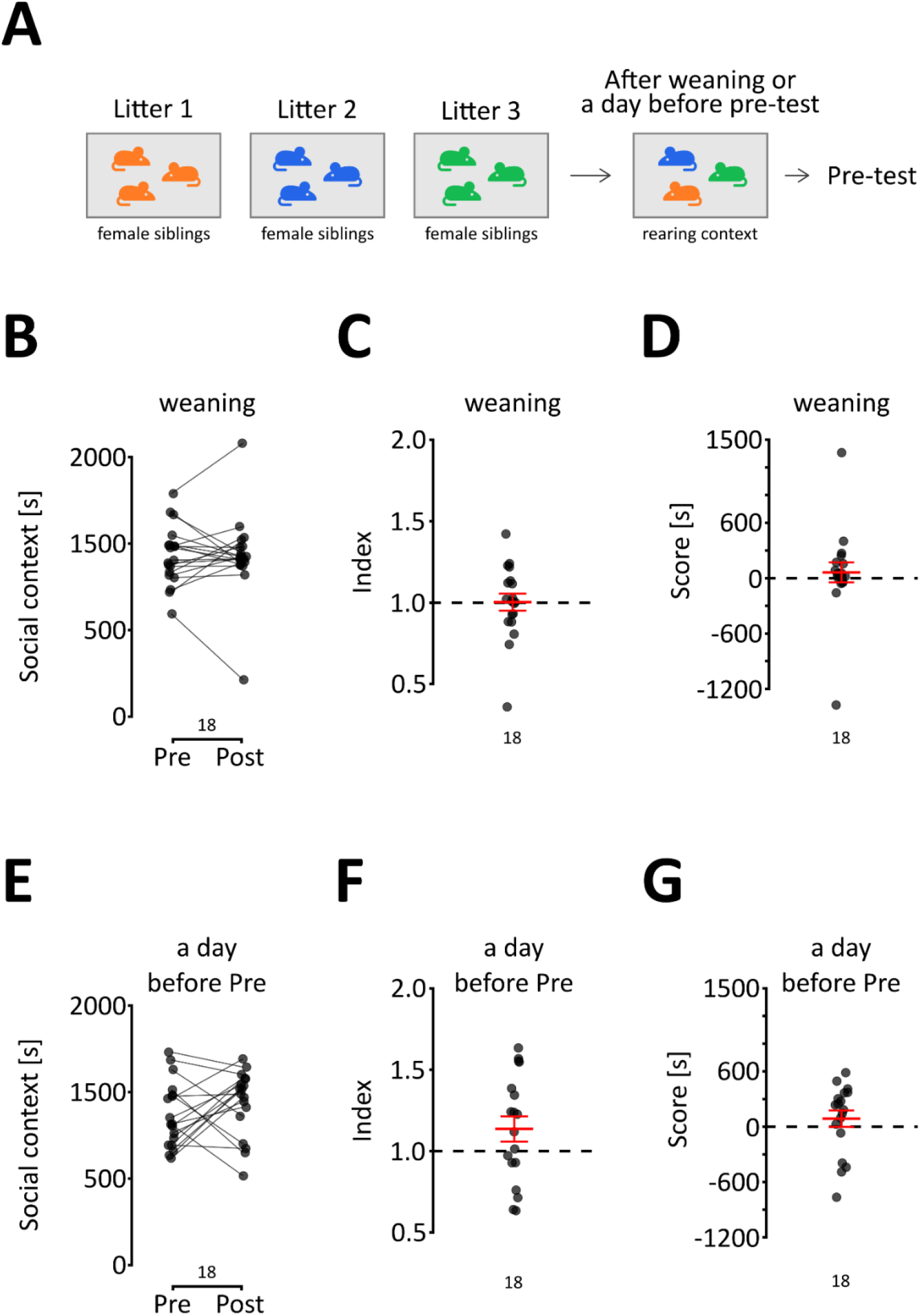
The effect of familiarity on social reward. (A, B and C) The change in time spent in the first context between pre- and post-test, the index and the score, respectively, in female mice that were brought together right after weaning. The mean and s.e.m. values are shown in red. (D, E and F) These graphs are the equivalents for female mice that were brought together one day before the pre-test. The mean and s.e.m. values are shown in red.

## Discussion

We show that in adult C57BL/6J female mice, contact with same-age female siblings is rewarding. Our results extend previous observations in juvenile mice^1,5^ and show that the protocol length and the type of context used have a critical role in the result of the sCPP task. Thus, we provide a behavioral method to study the mechanisms underlying the rewarding effects of nonreproductive social interaction in adult mice, which could be useful in models of human neuropsychiatric disorders that involve impaired social behaviors. Moreover, the revised procedure greatly facilitates the use of sCPP with genetically modified mice, where the age restrictions made the test impractical.

A critical element of the sCPP procedure is the length of the conditioning sessions. Here, we used 24-h conditioning periods for the social and isolation contexts, the same as several recent reports^1,3–5^. Protocols with short, 15-30-minute conditioning sessions have also been described, where one of the contexts was paired to interaction with an unfamiliar^11,12^ or a familiar^13^ juvenile or an age- and weight-matched partner. According to these reports, the rewarding effects of a brief interaction were also dependent on age and familiarity, and in contrast to this report, contact with unfamiliar mice resulted in significant place preference^11,12^. The effects of the brief contact with a familiar conspecific were reported to be strain specific; CD1 female mice showed robust sCPP, while C57BL/6J did not^13^. There are essential differences in the approaches, and we would like to note that the long conditioning periods exclude potential effects of novelty, sociability or social memory, while short conditioning sessions emphasize them and could be argued to share similarities to the three-chamber social approach task^14^.

We focused on female mice because our aim was to study the rewarding effects of amicable (as opposed to hostile) social interactions. As already noted, *Mus musculus* females show cooperative parenting behaviors^8^. Conversely, adult male mice in natural and seminatural environments show territorial behavior^15^. Under standard laboratory conditions, male mice also engage in fights, even with familiar partners. Fights are more frequent after environmental changes^16^. The sCPP protocol requires daily changes in bedding^1,5,6^; thus, it is likely to induce aggressive behavior. Furthermore, an opportunity for aggressive encounters has been consistently shown to induce place preference in male mice^9^. Therefore, if familiar male mice of the C57BL/6J strain engage both in hostile and affiliative social interactions^16^, it is not possible to distinguish which type of interactions are responsible for the conditioning effect. Hence, we believe that the neural basis of the rewarding effects of amicable social interactions between adult animals should be studied using female mice.

The observation that sCPP is absent in 14-week-old female C57BL/6 mice when mice are exposed to the social and isolation contexts only once was replicated (the two-day protocol^5^). It could be argued that the necessity to extend the conditioning to observe rewarding effects in adults proves that the effects of social contact are weak and thus difficult to observe in the sCPP task. However, we find such interpretation to be too speculative. First, the sCPP results should probably be considered qualitative rather than quantitative. Moreover, it would be difficult to distinguish the lack of rewarding properties of the stimulus from reduced salience or a slower learning rate. Additionally, it could be hypothesized that the isolation context conditioning plays a much stronger role in juvenile mice, as was previously noted^2^. Therefore, as a more plausible explanation, we found the effect of the length of conditioning reflects a difference in the underlying mechanism in adult vs. juvenile mice, although, admittedly this remains speculation without further evidence.

A particularly interesting observation is that the rewarding effects of social contact in adult female mice were only observed among siblings that were housed together from birth. It could argued that these observations show that social contact in adult female mice is rewarding only under very specific circumstances, or, alternatively that in adult animals the reward is proportional to familiarity, and might be too weak to be detected by sCPP. Either interpretation would be in line with the general notion that virtually all aspects of mammalian social behavior are influenced by familiarity between interaction partners^17–25^. In fact, in line with our observations, the communal nesting behavior was observed to be most likely in highly familiar animals^21^. Therefore, we were surprised that we could not find previously published reports that have directly assessed the influence of familiarity on the rewarding effects of social interactions in mice.

Two opposing hypotheses concerning the effect of familiarity on social reward can be derived from the literature. Cann and coauthors hypothesized that contact with unfamiliar individuals could be more rewarding for mice^2^. This hypothesis was based on their own observation of unstable conditioning effects elicited by social contact with familiar mice^2^ and more robust results acquired with a protocol using an unfamiliar individual as a stimulus animal^11^. However, two other lines of observations suggest that social contact with familiar individuals should be more rewarding than with unfamiliar individuals. First, both male and female mice engage in affiliative social interactions more frequently with familiar than unfamiliar conspecifics^20,26–28^. The second observation concerns the reproductive success of communally rearing pairs of female mice of different degrees of familiarity and relatedness; offspring survival probability is higher for females that were reared together than for females grouped as young adults, irrespective of their relatedness status^21^. The results presented here are in line with these observations. We have shown that social contact with siblings reared in the same cage is rewarding for adult female mice, while social contact between conspecifics familiarized after weaning does not have rewarding properties. However, no conclusions concerning the influence of genetic relatedness on social reward could be derived from our study, as it was performed on an inbred mouse strain (C57BL/6). It has been shown that mice recognize kin by detecting the major urinary proteins (MUPs)^29^ (but see^30^), and the genetic diversity of MUPs is very low among common laboratory mouse strains, even outbred^31^. Hence, mice from inbred strains likely perceive all members of their strain as kin. We hypothesize that the results of our study could be explained by familiarization with sibling cues at a very early age or even during the prenatal period. Future research should assess the impact of genetic relatedness on social reward in mice.

## Acknowledgments

This work was supported by grant OPUS UMO-2016/21/B/NZ4/00198 from the National Science Centre Poland and the statutory funds of the Maj Institute of Pharmacology of the Polish Academy of Sciences.

## Methods

### Animals

Adult female C57BL6 mice were bred at the Animal Facility of the Maj Institute of Pharmacology, Polish Academy of Sciences. The exact age and sex of the animals used for the initial screening of bedding sets are shown in Table S2. The groups of animals used in the sCPP experiments are described in Table S3. All behavioral procedures were approved and monitored by the II Local Bioethics Committee in Krakow (permit numbers 224/2016, 35/2019, 266/2020, 38/2021, 67/2020) and performed in accordance with the Directive 2010/63/EU of the European Parliament and of the Council of 22 September 2010 on the protection of animals used for scientific purposes. The reporting in the manuscript follows the recommendations in the ARRIVE guidelines.

Before the experiment, animals were group housed (2-6 per cage) on aspen bedding in standard type II L cages. One of two similar brands of aspen bedding was used (Table S4). Gnawing blocks and bedding material were always present. Standard rodent chow and water were available *ad libitum.* The environmental conditions were as follows: temperature (22 ± 2 °C), humidity (app. 40-60%), and a 12/12-h light-dark cycle with lights on at 7 AM CET/CEST. Pre- and post-tests were conducted during the light phase.

Animals were weaned at the age of 27-28 days, and female offspring were placed in a new cage. For the basic version of the protocol (Figures 1–3), mice were housed with their sisters since weaning throughout the experiment, except in the experiments where the effect of familiarity was tested. In the first of the two experiments assessing the effect of familiarity (Figure 4, upper panel), mice were housed after weaning (4 weeks of age) and tested in nonsibling groups. In the second experiment (Figure 4, lower panel), animals were housed together with their sisters from weaning until the day before the pre-test (at approximately 12 weeks of age). On the day before the pre-test, they were mixed with mice from different litters and housed in the new groups throughout the experiment. In both experiments, mice were mixed in such a way that only one animal from a given litter was present in the cage. The animals were handled for 4-5 days before the beginning of the experimental procedures.

### Conditioning contexts

Four sets of conditioning contexts were tested (Table S1, Figure 1). Each set consisted of two distinct environmental contexts (context A and context B) that differed in bedding type (Table S4) and gnawing block size and shape (Table S5). The test for the context preference was performed in a custom-made opaque plastic cage (30 × 30 × 30 cm) divided into two identical compartments by a transparent plastic wall, with a 5 × 5-cm opening at the base. Each compartment contained one type of context. Mice were allowed to freely explore the cage for 30 minutes. Two animals were tested simultaneously in cages placed adjacently. Behavior was recorded using a camera placed above the cages, and the amount of time spent in each compartment was measured automatically using EthoVision XT 15 software (Noldus, The Netherlands). The experimental room was dimly lit (5-10 lux at the bottom of the cage). Additional illumination was provided with near-infrared LED lights.

### Social conditioned place preference test (sCPP)

The procedure was performed as previously described^10^. The test consisted of three phases: the pre-test, conditioning phase, and post-test (Figure 1A). The pre-test and post-test were performed exactly like the test for the context preference (see above). After the pre-test, animals were returned to their home cage for approximately 24 h. Then, mice were assigned to undergo social conditioning (housing with cage mates) for 24 h in one of the contexts used in the pre-test followed by 24 h of isolation conditioning (single housing) in the other context. For an unbiased design, the social context was randomly assigned in such a way that approximately half of the animals received social conditioning in context A and half in context B. The conditioning phase lasted 6 days (3 days in each context, alternating every 24 h, Figures 1, 2, 4) or 2 days (1 day in each context, Figure 3). After conditioning, the post-test was performed.

In the case of the experiment where effects independent of social contact were assessed, animals were housed together during all 6 conditioning phases (in both contexts).

To determine if mice developed social preference, we used three criteria: 1) pre-test-post-test comparison of the time spent in the social context, 2) social score – the difference in time spent in the social minus isolation context during the post-test (comparted to chance value, i.e., 0), 3) social index – the ratio of time spent in the social context during the post-test and time spent in social context during the pre-test (comparted to chance value, i.e., 1).

### Data analysis

All results are reported in accordance with the ARRIVE 2.0 guidelines.

Animals that spent more than 70% of the pre-test time in one of the contexts were excluded from the analysis. In the instances when the number of animals conditioned on different bedding types were not equal (i.e., in the experiments assessing social context-independent effects and the influences of age and familiarity), the number of animals for each type of bedding was equalized by randomly removing the necessary number of cases from the larger group. The R script used for trimming the larger group and the datasets before and after trimming are available from: https://doi.org/10.5281/zenodo.6347482.

The effects of bedding on time spent in the compartments were assessed using analysis of variance followed *post hoc* by Bonferroni corrected t test, two-sample t test or one sample t test as appropriate.

## Author contributions

ZH and JRP planned the research; ZH, MC, KM, MK, ŁS, and MK-J performed the experiments; ZH, MC, KM, MK, and JRP analyzed the data, and ZH and JRP wrote the manuscript with help from all the authors.

## Data availability statement

All data are available at https://doi.org/10.5281/zenodo.6347482. Raw video recordings of sCPP trials will be made available on request.

